# Language impairments in ASD resulting from a failed domestication of the human brain

**DOI:** 10.1101/046037

**Authors:** Antonio Benítez-Burraco, Wanda Lattanzi, Elliot Murphy

## Abstract

Autism spectrum disorders (ASD) are pervasive neurodevelopmental disorders entailing social and cognitive deficits, including marked problems with language. Numerous genes have been associated with ASD, but it is unclear how language deficits arise from gene mutation or dysregulation. It is also unclear why ASD shows such high prevalence within human populations. Interestingly, the emergence of a modern faculty of language has been hypothesised to be linked to changes in the human brain/skull, but also to the process of self-domestication of the human species. It is our intention to show that people with ASD exhibit less marked domesticated traits at the morphological, physiological, and behavioural levels. We also discuss many ASD candidates represented among the genes known to be involved in the domestication syndrome (the constellation of traits exhibited by domesticated mammals, which seemingly results from the hypofunction of the neural crest) and among the set of genes involved in language function closely connected to them. Moreover, many of these genes show altered expression profiles in the brain of autists. In addition, some candidates for domestication and language-readiness show the same expression profile in people with ASD and chimps in different brain areas involved in language processing. Similarities regarding the brain oscillatory behaviour of these areas can be expected too. We conclude that ASD may represent an abnormal ontogenetic itinerary for the human faculty of language resulting in part from changes in genes important for the domestication syndrome and, ultimately, from the normal functioning of the neural crest.

## 1. Introduction

Autism spectrum disorders (ASD) are pervasive neurodevelopmental conditions characterised by several and severe cognitive and social deficits, including language and communication problems, repetitive and stereotypical behaviour, and problems with social interaction (Bailey et al. 1996). In DSM-V, language deficits are no longer explicitly postulated as a central feature of ASD because they are subsumed in its distinctive communication problems. Nevertheless, it is clear that ASD entails a typical language profile and language developmental path (reviewed in Benítez-Burraco and Murphy 2016; see also Tager-Flusberg et al. 2005, Tager-Flusberg 2006, Eigisti et al. 2007, Bourguignon et al. 2012). Because of the masking effect of a variable IQ, and the variable degree of functionality exhibited by ASD patients, it is difficult to hypothesise a core language deficit in this condition. The impairment of the oromotor function has been claimed to account for expressive language problems in some autistic subjects (Belmonte et al. 2013). Comprehension problems seemingly result from other underlying deficit(s), including a reduced effect of semantic priming (Preissler 2008), problems with phonological processing (Lindgren et al. 2009), or impairment of procedural memory (Walenski et al. 2006).

At the neural level, ASD entails atypical development, wiring and interconnection of areas involved in language processing (Stefanatos and Baron 2011, Bourguignon et al. 2012). Not surprisingly, functional differences in language processing tasks of ASD compared with unaffected subjects have been attested as well (Courchesne and Pierce 2005, Scott-Van Zeeland et al. 2010a, b). For instance, microstructural anomalies and reduced lateralization patterns have been observed in the arcuate fasciculus of ASD patients (Fletcher et al. 2010), suggesting that a constraint on the integrative processes during development may contribute to language impairment in this condition (Schipul et al. 2011). We also wish to highlight both increased and decreased intra‐ and inter-hemispheric connectivity (Hahamy et al. 2015), and abnormal responses to linguistic stimuli (reviewed in Stefanatos and Baron 2011: 259-262). Intriguingly, the ASD phenotype is characterised by increased intrinsic functional connectivity during the first years of life (the time window where language is acquired) and reduced connectivity in adolescent and adult states (Uddin et al. 2013).

In spite of this growing body of neurobiological data, a comprehensive view of language processing in the ASD brain is still lacking. Specifically, ASD studies need to move beyond simplistic models of language processing and focus instead on how collections of brain areas jointly engaged in specific, impaired cognitive operations (see Fedorenko and Thompson-Schill 2014 for a general discussion). This is a real challenge, provided that abnormal brain profiles are not expected to easily map on to anomalous categories or computations of linguistic theories (see Poeppel 2012 and Murphy 2016 for discussion). We have recently proposed a translational theory of language deficits in ASD as amounting to abnormal patterns of brain rhythms (Benítez-Burraco and Murphy 2016); although a clarification and empirical validation of this hypothesis is still pending.

Finally, we wish to emphasize that ASD has been associated with sequence variants, copy number variation (CNVs), and/or changes in the expression patterns of an extensive number of genes (Geschwind and State 2015). Despite the remarkable genetic heterogeneity, it is noteworthy that all these genes tend to converge on specific pathways and neural mechanisms, functionally relevant in this condition and expected to account for its associated deficits (Jeremy Willsey and State 2014). Specifically, several candidates for language impairment in ASD have been proposed, including *MET, CTTNBP2, EN2, NBEA, HRAS*, and *PTEN* (Campbell et al. 2006, Cheung et al. 2001, Benayed et al. 2005, Castermans et al. 2003, Comings et al. 1996, Naqvi et al. 2000). Nonetheless, the gap between genes and language deficits in ASD still remains open (see Jeste and Geschwind 2014 for a general discussion, and Benítez-Burraco and Murphy 2016 for a specific discussion on candidates for language dysfunction in ASD).

The aim of this paper is to contribute to the bridging of the gap between the genetic backdrop and language deficits observed in ASD. To this end, we will primarily focus on language evolution. There exists a strong, deep link between evolution and (abnormal) development. Recently-evolved neural networks seem to be more sensitive to damage because of their lower levels of resilience (Toro et al. 2010). As a consequence, aspects of brain development and function that are preferably impaired in modern populations are expected to be involved in recently evolved, human-specific cognitive abilities. Some comprehensive accounts of the human condition set against the cognitive profiles of other primates have been recently put forth (Seed and Tomasello 2010, Platt et al. 2016). Comparative genomics also provides valuable information about the sources of the observed differences and similarities in the human genome (Rogers and Gibbs 2014, Franchini and Pollard 2015). Likewise, we are beginning to achieve an advanced understanding of the genetic changes that occurred after our split from extinct hominins (Pääbo 2014, Zhou et al. 2015). We expect that the same factors that prompted the transition from an ape-like cognition to our specific mode of cognition are involved in the aetiology of cognitive disorders involving language deficits and, particularly, of ASD (see Benítez-Burraco 2016 for a general discussion).

In what follows, the focus is placed on one aspect of this evolutionary process: the self-domestication of the human species. At present, we have a decent understanding of how our language-readiness (that is, our species-specific ability to learn and use language) may have evolved. Accordingly, among the changes brought about by human evolution, one very relevant aspect is the ability to transcend (better than other species) the signature limits of core knowledge systems and thus go beyond modular boundaries (Mithen 1996, Spelke 2003, Carruthers 2006, Hauser 2009, Boeckx 2011, Wynn and Coolidge 2011). As hypothesised in Boeckx and Benítez-Burraco (2014a), our language-readiness boils down to this enhanced cognitive ability, but also to its embedding inside cognitive systems responsible for interpretation (*thought*) and externalization (*speech*). This language-readiness was seemingly brought about by specific changes in the skull/brain developmental path (resulting in a more globular brain), which entailed new patterns of long-distance connections among distributed neurons and, ultimately, new patterns of brain rhythmicity, including an adequate degree and pattern of cortical inhibition. Interestingly, brain rhythms are heritable components of brain function (Linkenkaer-Hansen et al. 2007, Hall et al. 2011) and have been linked to computational primitives of language (Murphy 2015a, b, 2016), allowing for a good explanation (and not just a description) of linguistic computation (and of language deficits) at the brain level, and specifically, for a satisfactory mapping of language deficits to neural dysfunction and its genetic basis in ASD (Benítez-Burraco and Murphy 2016). We have found many candidates for ASD among the genes known to be involved in the emergence of language-readiness (Benítez-Burraco and Boeckx 2015).

At the same time, the emergence of modern-like languages (and perhaps of core aspects of language too) was seemingly favoured by changes in the cultural niche of our ancestors. The archaeological record shows that cognitive modernity (encompassing language-readiness) did not automatically entail behavioural modernity (seemingly resulting from using fully-fledged languages), which only appeared long after the emergence of anatomically-modern humans (AMHs) together with changes in human cultural dynamics. Current linguistic research has shown that aspects of linguistic complexity (including core aspects of grammatical knowledge) correlate with aspects of social complexity (Lupyan and Dale 2010, Wray and Grace 2007). Moreover, core properties of human languages (like duality of patterning) can develop in response to environmental pressure, as research into emergent sign languages has nicely illustrated, implying that they cannot be regarded as part of the biological endowment (see Benítez-Burraco 2016b for discussion). Importantly, language acquisition by the child demands a prolonged socialization window that enables her to receive the proper amount of triggering stimuli and to interact with other conspecifics. All this means that the intrinsic cognitive machinery may be not enough for granting the acquisition of a successful tool for linguistic cognition and that the environment has to be of the right kind too (see Sterelny 2011 on behavioural modernity set against cognitive modernity).

It has been hypothesised that the social conditions (or the *cultural niche*) that facilitated the enhancement of linguistic structure through a cultural mechanism were brought about by a process of human self-domestication (see Thomas 2014 for details, and Hare and Tomasello 2005 and Deacon 2009 on relaxed selective pressures resulting from self-domestication as explanations of the emergence of key aspects of behavioral modernity). Different factors may have contributed to human self-domestication, from adaptation to the human-made environment to selection against aggression to sexual selection. We have hypothesised (Benítez-Burraco et al. submitted) that the very changes that brought about our globular skull/brain and our language-readiness may have also fuelled the emergence of a (self-domesticated) phenotype in the human species. Accordingly, we have found numerous links between the candidates for globularization and language-readiness, and genes important for the development and function of neural crest cells (NCC). Indeed, the hypofunction of the neural crest (NC) has been claimed to account for the constellation of distinctive traits observed in domestic mammals (the “domestication syndrome”) (Wilkins et al. 2014).

Because of the deep link between evolution and development, we expect that examining the signatures of the domesticated phenotype in people with ASD contributes to a better understanding of aetiology of ASD, and specifically, of language deficits in this condition. In a recent paper Reser (2014) found similarities between autism and species of solitary mammals. Although the focus was put on behaviour, the author suggests that future research will benefit from investigating the neurobiological, genetic and epigenetic causes of these similarities. Here we try to push research in this direction. The paper is structured as follows. First, we provide a general account of the domesticated traits that are absent or attenuated in ASD. Then we move to the genes and focus on candidates for ASD that are found among the set of genes involved in the domestication syndrome and the evolution of language-readiness, as characterised in Benítez-Burraco et al. (submitted), showing that they exhibit a distinctive expression profile in the brain of autists. Finally, we compare the ASD phenotype with wild primates, focusing on the expression profile of these genes, but also on oscillatory signatures of areas important for language processing, considering that language impairment in ASD can be interpreted as an ‘oscillopathic’ condition (see Benítez-Burraco and Murphy 2016). We will conclude that ASD (and language deficits in ASD) can be viewed as an abnormal ontogenetic itinerary for the human faculty of language, resulting in part from changes in genes important for the domestication syndrome and seemingly from changes in the normal functioning of the NC.

## 2. Domestic traits in the ASD phenotype

Wilkins et al. (2014) provide a comprehensive summary of traits known to be modified in domesticated mammals, many of them concerning the cranial region. These include changes in ear size and shape, changes in the orofacial area (including shorter snouts and smaller jaws), changes in dentition (particularly, smaller teeth), and a reduced brain capacity (specifically, of components of the forebrain such as the amygdala or parts of the limbic system). Other distinctive traits commonly found in domesticated strains are depigmentation, neoteny, shorter reproductive cycles, and increased docility, which is thought to result from adrenal size reduction and adrenal hypofunction as well as from reduced levels of stress hormones (including adrenocorticoids, adrenocorticotropic hormone, cortisol, and corticosterone). This delayed adrenal maturation also involves a hypofunction of the sympathetic nervous system and an increase of the duration of the immaturity of the hypothalamic-pituitary-adrenal system (the HPA axis), which provides the animal with a longer socialization window. According to Wilkins et al. (2014), the multiple phenotypic traits that characterize the domestication syndrome emerge as unselected by-products from a developmental reduction in NCC inputs, resulting from selection for tameness. Interestingly, compared to extinct hominins, AMHs exhibit a number of domesticated traits, including reduced brains (at least during the last 50,000 years), changes in dentition, reduction of aggressiveness, and retention of juvenile characteristics (see Thomas 2014 for details). Intriguingly, most of these features are generally attenuated in ASD (Figure 1).

**Figure 1.**
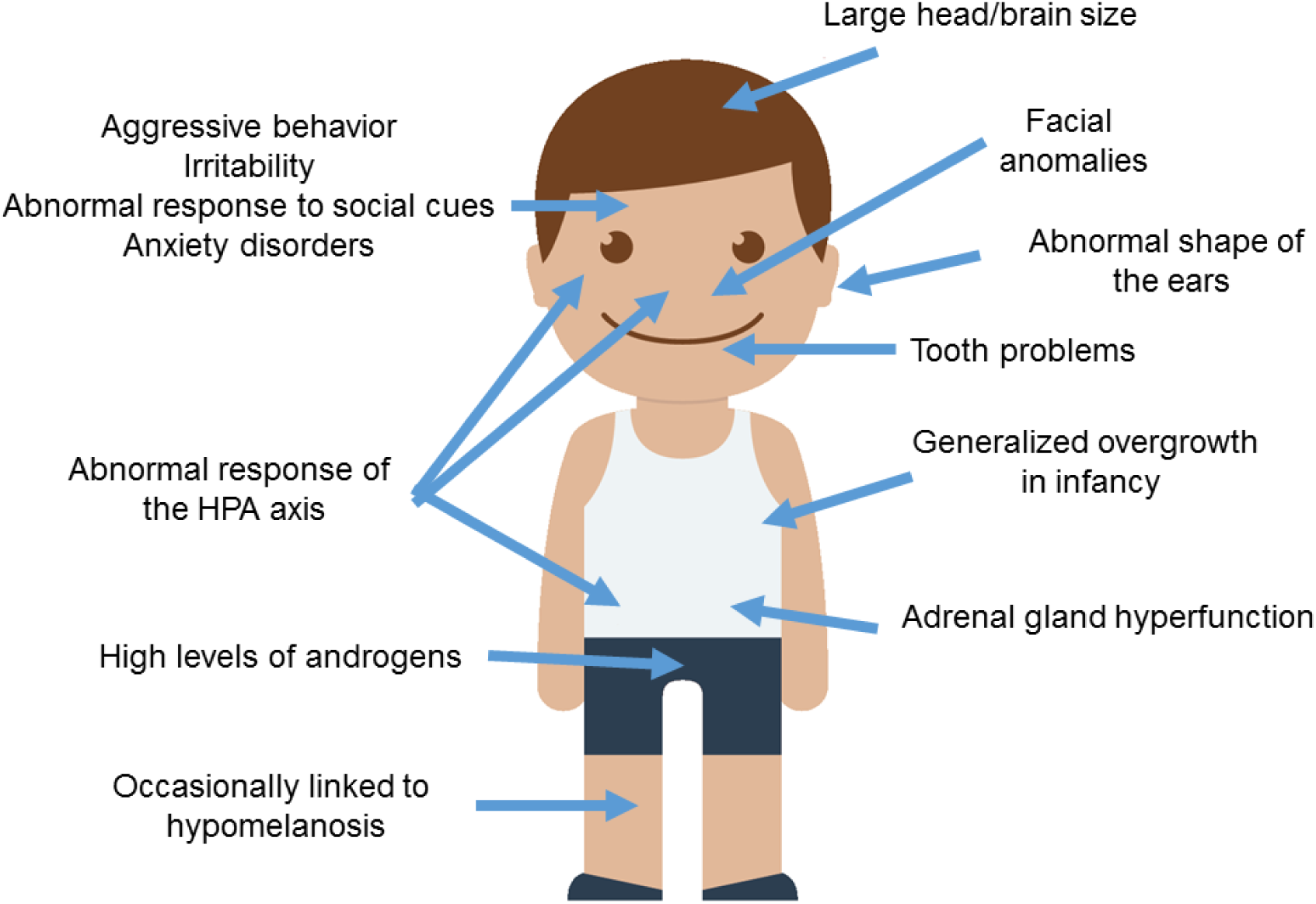
Anomalous presentation of domesticated traits in people with ASD. The image of the child is from Iconfinder (https://www.iconfmder.com/icons/525448/boy_child_kid_male_man_person_white_icon).

To begin with, ASD subjects show significant differences with healthy controls regarding minor physical anomalies, particularly in the craniofacial region (assumed to result from deviations during foetal development and suggested to constitute external markers of atypical brain growth) (Tripi et al. 2008, Manouilenko et al. 2014). Specifically, in adults the abnormal shape of the ears is robustly associated with autistic traits, with higher scores correlating with poorer functioning (Manouilenko et al. 2014). Regarding the changes in the orofacial region, prepubertal boys with ASD show significant differences in facial morphology compared to typically developing (TD) boys (Aldridge et al. 2011). This distinctive facial phenotype is more pronounced in subjects with severe symptoms, significant cognitive impairment, and language regression (Obafemi-Ajayi et al. 2015). Concerning tooth peculiarities, children with ASD show greater abnormalities in dentition, including missing teeth, diastemas, or reverse overjets (Luppanapornlarp et al. 2010). With respect to brain size, head circumference is significantly larger in people with ASD, with nearly 15% suffering from macrocephaly. Higher brain volumes correlate with lower functioning abilities; indeed nearly 9% ASD individuals exhibit brain overgrowth (Sacco et al. 2015). It is worth noticing that higher head circumference and brain size values are observed only during early childhood (Fukumoto et al. 2008, Courchesne et al. 2011, although see Raznahan et al. 2013), particularly when ASD is presented with regression (Nordahl et al. 2011). Typically, early brain overgrowth is followed by a decrease in structural volumes (Courchesne et al. 2011). Although brain overgrowth may result from a dysregulation of the overall systemic growth (see below), it is thought to impact on cognition. This is believed to occur as a result of the reduced networking efficiency among widespread regions of the cortex, due to the increased long-distance connections (Lewis et al. 2013). Specifically, people with ASD show increased volumes of the amygdala (Mosconi et al. 2009, Murphy et al. 2012), which correlate with the severity of their social and communication impairments (Schumman et al. 2009). In the TD population, higher amygdala volumes are associated with poorer language abilities in infancy (Ortiz-Mantilla et al. 2010).

Regarding the behavioural traits associated with the domestication syndrome, we wish to highlight that aggressive behaviours are frequent in children with ASD (with about 25% of them having scores in the clinical range), and correlate with lower cognitive outcomes (Hill et al. 2014). Children with ASD display more reactive than proactive aggression attitudes (Farmer et al. 2015). Likewise, irritability is also commonly observed in affected individuals (Mikita et al. 2015). Additionally, ASD is commonly found to be comorbid with generalised anxiety disorder (Hollocks et al. 2014, Bitsika et al. 2015). Several studies have been carried out to learn more about the physiological basis of this anomalous response to the social environment. Interestingly, higher serum cortisol responses are usually found in children with ASD, particularly after stressor stimulation, when prolonged duration and recovery of cortisol elevation is also observed (Spratt et al. 2012). Moreover, children with ASD show a distinctive diurnal rhythm of cortisol compared to their TD peers; this involves elevated cortisol levels at the end of the day and dampened linear decline across the day in some children (Tomarken et al. 2015). Dysregulation of the diurnal rhythm as a whole has been found in low functioning ASD (Taylor and Corbett 2014). Also, anxiety symptoms correlate with high cortisol levels in ASD paediatric patients (Bitsika et al. 2015). Plasma levels of adrenocorticotropic hormone are also significantly higher in children with ASD, and correlate positively with the severity of the symptoms (Hamza et al. 2010). The HPA axis in ASD responds in a more sluggish way to physiological or physical manipulation. Accordingly, Taylor and Corbett (2014) found hyper-responsiveness of the HPA axis when unpleasant stimuli or relatively benign social situations are involved, whereas they observed hypo-responsiveness in conditions involving social evaluative threat. On the whole, the HPA axis may be more reactive to stress in social anxiety disorder and ASD (Spratt et al. 2012, Jacobson 2014). Because children with autism and anxiety disorders show a blunted cortisol response to psychosocial stress, and given that reduced cortisol responsiveness is significantly related to increased anxiety symptoms, Hollocks et al. (2014) suggested that a non-adaptive physiological response to psychosocial stress may exist in ASD.

Finally, it is worth considering some other traits commonly observed in domesticated mammals: neoteny, alterations of reproductive cycles, and pigmentation changes. Regarding neotenic features, it is noteworthy that children with ASD exhibit an early generalized overgrowth (Van Daalen et al. 2007, Fukumoto et al. 2008, Chawarska et al. 2011). Typically, boys with ASD show increased body size at birth and during infancy, with postnatal overgrowth correlating with lower adaptive functioning, greater severity of social deficits, and poorer verbal skills (Chawarska et al. 2011, Campbell et al. 2014). Interestingly, higher levels of androgens are found in children and adolescents with ASD. This correlates with the severity of autistic traits and might account for the precocious puberty also reported in this condition (El-Baz et al. 2014). These findings emphasize the role of elevated pre‐ and postnatal testosterone levels in the liability for ASD (see Hauth et al. 2014). Testosterone significantly affects brain development, particularly targeting the hypothalamus, the amygdala and the hippocampus, impacting on aspects of memory consolidation (Filová et al. 2013). High perinatal testosterone concentration negatively correlates with early vocabulary development in TD boys (Hollier et al. 2013). Interestingly, children with elevated androgen levels due to congenital adrenal hyperplasia show atypical patterns of brain asymmetry in the perisylvian areas, and language/learning disabilities (Plante et al. 1996). Less data on reproductive functions in females is available, due to the lower prevalence of ASD among women. Nevertheless, women with ASD reported significantly more irregular menstrual cycles and dysmenorrhea (Ingudomnukul et al. 2007, Hamilton et al. 2011). Likewise, an increase in premenstrual syndrome has been observed in women with ASD (Obaydi and Puri 2008, Hamilton et al. 2011), who are more likely to exhibit behavioural issues related to the onset of periods (Burke et al. 2009). In addition, delayed age of menarche seems to correlate with the severity of autistic traits (Hergüner and Hergüner 2016). These findings lend support to the androgen theory of ASD, according to which elevated levels of testosterone during foetal development may contribute to the development of ASD. Finally, concerning changes in pigmentation, it is of interest that hypomelanotic diseases usually entail autistic symptoms, as is commonly observed in hypomelanosis of Ito (OMIM#300337; Akefeldt and Gillberg 1991, Von Aster et al. 1997, Gómez-Lado et al. 2004). It has been hypothesised that the comorbidity between hypomelanosis and ASD may result from a deficiency in vitamin D (Eyles 2010, Bakare et al. 2011). In fact, core symptoms of ASD improve after vitamin D supplementation (Jia et al. 2015). Interestingly, core candidates for the globularization of the AMH skull/brain and the evolution of language-readiness are involved in vitamin D homeostasis and function (see Benítez-Burraco and Boeckx 2015 for details).

As noted above, regardless of the different selectionist scenarios that may account for the traits commonly found in domesticated mammals, a role for NC hypofunction during embryonic development has been proposed (see Wilkins et al. 2014 for details). No comprehensive view of the role (if any) of the NC in the aetiopathogenesis of ASD has been provided to date. Still, it is important to note that neurocristopathies (that is, conditions resulting from NC defects) commonly involve autistic features. For instance, in CHARGE syndrome (OMIM#214800) autistic traits coexist with developmental abnormalities affecting endocrine, reproductive, urinary and digestive systems, along with skeletal and craniofacial features (Fernell et al. 1999). Given this background, we will now examine whether candidates for ASD are overrepresented among the genes believed to play a central role in NC development and function, with a special emphasis on those that interact with genes important for the globularization of the AMH skull/brain.

## 3. ASD and the genetics of the domestication syndrome

In order to improve our characterization of the domesticated traits in ASD, it is of interest to assess whether candidate genes for this condition (with a particular emphasis on language disabilities) are overrepresented among, or are functionally related to, candidates for domestication. We have relied on an extended list of candidates, which includes the core set of genes proposed by Wilkins et al (2014), plus a subset of the genes involved in the globularization of the AMH skull/brain and the emergence of language-readiness that are functionally related to them, through direct interaction, and/or that play a role in the development and function of the NC (see Benítez-Burraco et al. submitted for details). Our list also comprises NC-related genes known to play a key role in craniofacial development and/or disorders. As noted above, most of the domesticated traits result from the modification of the cranial region and many of the ASD distinctive features concern the skull, face and brain. Moreover, as reasoned in Boeckx and Benítez-Burraco (2014a, b), we expect that our language-readiness resulted from changes in the development of the skull/brain, but also from the refinement of the externalization devices, specifically, the orofacial region: As also noted above, the impairment of oromotor function has been hypothesised to account for some language deficits in ASD. Table 1 provides a full list and a schematic characterization of these genes.

**Table 1.**
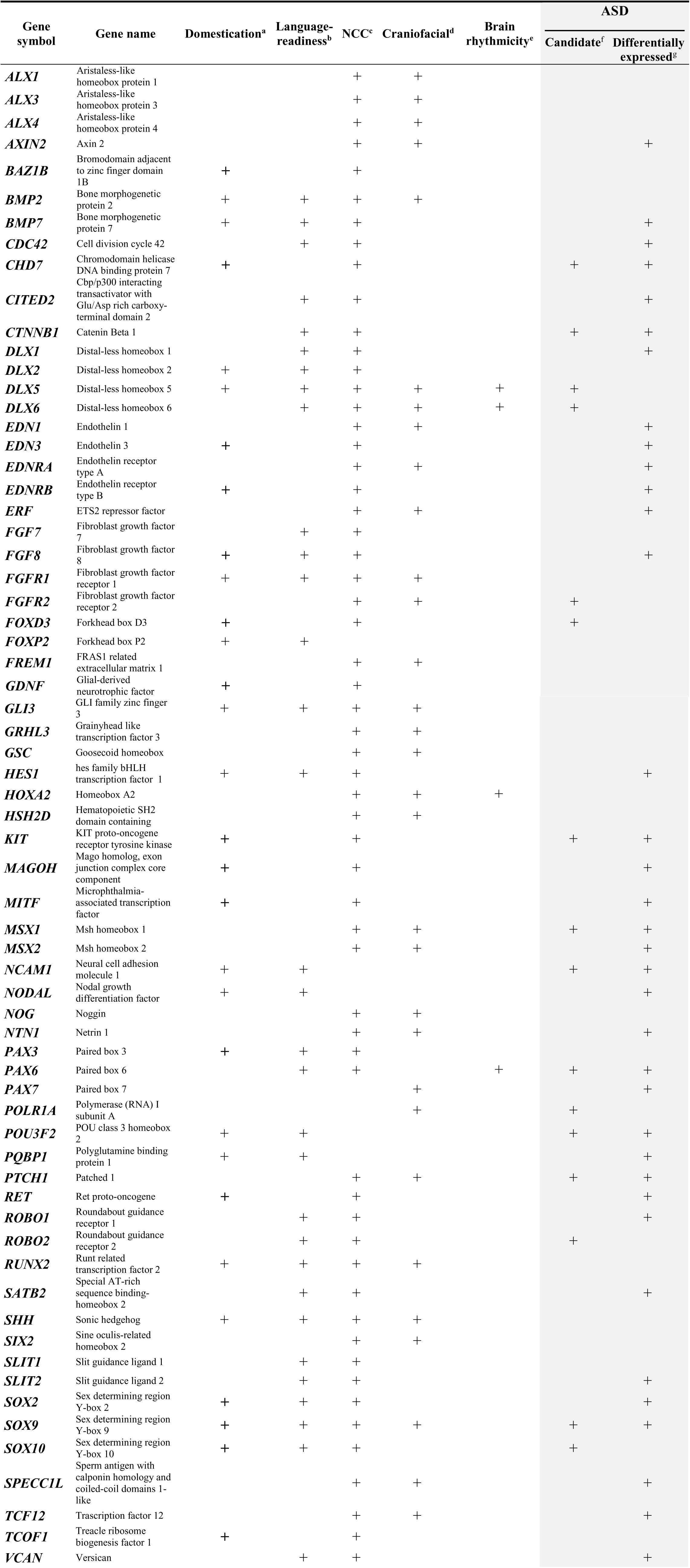
List of putative candidate genes for domestication and ASD. Notes: a. Core candidates for the “domestication syndrome” according to Wilkins et al. (2014) (bold italicized tags) plus language-readiness genes functionally interacting with them according to Benítez-Burraco et al. 2016 (regular tags). b. Genes highlighted as candidates for globularization of the AMH skull/brain and the emergence of language-readiness according to Boeckx and Benítez-Burraco 2014a, b and Benítez-Burraco and Boeckx 2015). c. Involved in neural crest (NC) development and function. d. Involved in craniofacial development and/or found mutated in craniofacial syndromes. e. Involved in brain oscillation and rhythmicity. f. Candidate for ASD as resulting from genomic studies (pathogenic SNPs, association studies, CNVs, functional studies, etc.). g. Differentially expressed in post-mortem brain tissues of ASD-versus-control individuals (see text for details).

When we tried to identify ASD-candidates among this extended list of genes via PubMed (http://www.ncbi.nlm.nih.gov/pubmed), we found out that nearly 25% of them have been suggested to play a role in the aetiopathogenesis of ASD. If we also consider genes that we found differentially expressed in post-mortem brain tissues isolated from patients (as discussed in the subsequent section), the percentage rises above 50%. Interestingly, some of these genes are thought to be involved in brain rhythmicity (see Table 1), plausibly contributing to the oscillopathic signature of the ASD brain during language processing.

We expect that the genes we highlight here are functionally interconnected and map on to specific pathways, signaling cascades, or aspects of brain development and function, of interest for language processing and the aetiopathology of ASD. In silico analyses offer promising insights. Accordingly, String 10 (www.string-db.org) predicts quite robust links between most of these genes (Figure 2). Likewise, ontology analyses by Panther (www.pantherdb.org) suggest that they might play biological functions important for ASD and be part of signaling pathways known to be impaired in this condition (Table 2).

**Figure 2.**
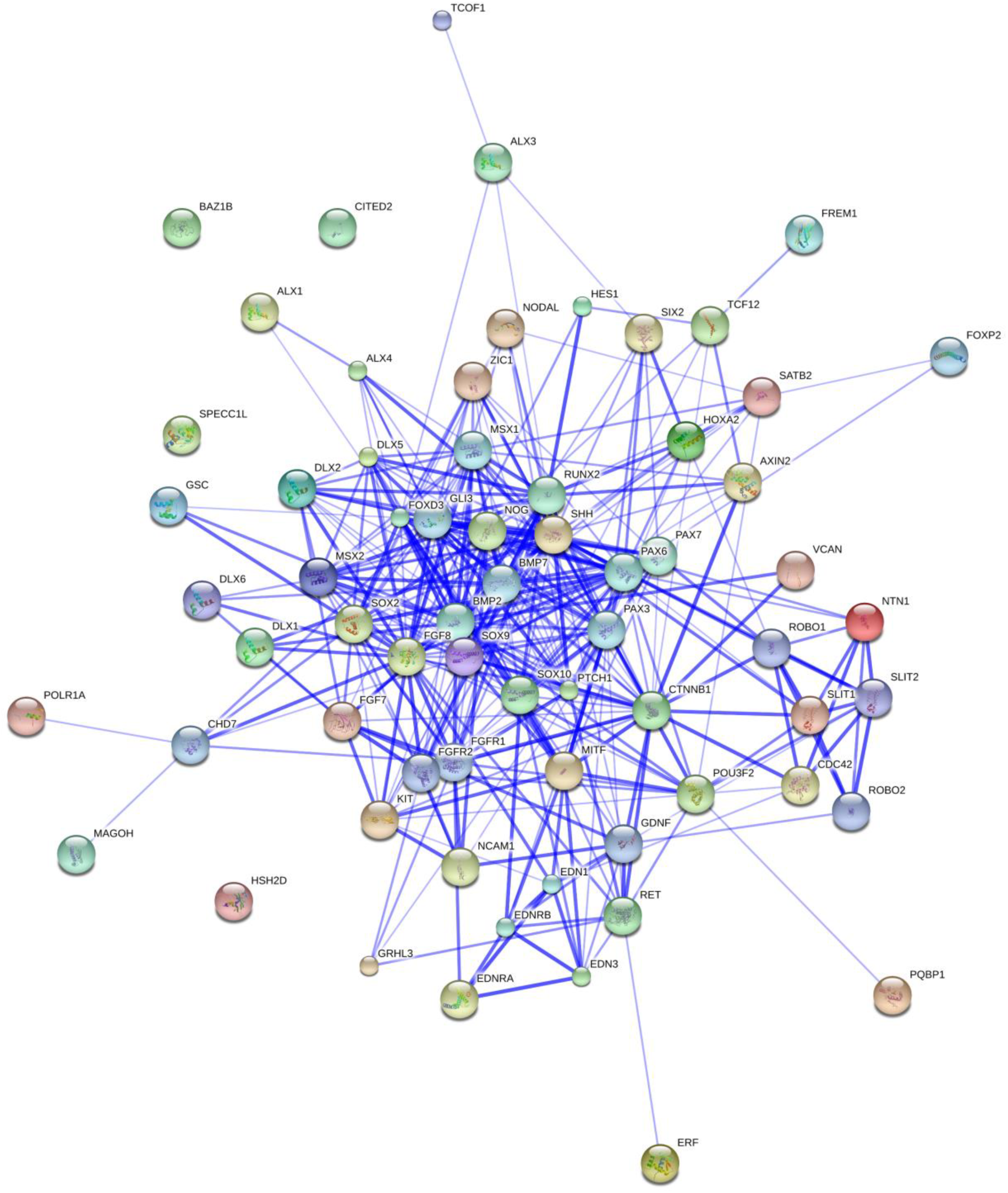
Functional links predicted by String 10 among candidates for domestication and ASD (Table 1). Stronger associations between proteins are represented by thicker lines. The medium confidence value was .0400 (a 40% probability that a predicted link exists between two enzymes in the same metabolic map in the KEGG database: http://www.genome.jp/kegg/pathway.html). String 10 predicts associations between proteins that derive from a limited set of databases: genomic context, high-throughput experiments, conserved coexpression, and the knowledge previously gained from text mining (Szklarczyk et al. 2015). This is why the figure does not represent a fully connected graph (evidence for additional links are provided in the main text). Importantly, the diagram only represents the potential connectivity between the involved proteins, which has to be mapped onto particular biochemical networks, signaling pathways, cellular properties, aspects of neuronal function, or cell-types of interest that can be confidently related to aspects of language development and function.

**Table 2.** GO classifications of candidates for domestication and ASD (Table 1), as provided by Panther (http://pantherdb.org). Numbers refer to percent of gene hit against total of process or pathway hits. Only the top ten functions after a Bonferroni correction have been included.

| **Biological process** |  | **Pathway** |  |
| --- | --- | --- | --- |
| metabolic process<br>(GO:0008152) | 21,60% | Axon guidance mediated by<br>Slit/Robo (P00008) | 12,20% |
| biological regulation<br>(GO:0065007) | 18,80% | TGF-beta signaling pathway<br>(P00052) | 10,20% |
| developmental process<br>(GO:0032502) | 17,00% | Endothelin signaling pathway<br>(P00019) | 8,20% |
| cellular process<br>(GO:0009987) | 13,80% | Gonadotropin releasing hormone<br>receptor pathway (P06664) | 8,20% |
| multicellular organismal<br>process (GO:0032501) | 11,50% | Wnt signaling pathway (P00057) | 8,20% |
| immune system process<br>(GO:0002376) | 4,10% | FGF signaling pathway (P00021) | 8,20% |
| apoptotic process<br>(GO:0006915) | 3,20% | Angiogenesis (P00005) | 6,10% |
| response to stimulus<br>(GO:0050896) | 3,20% | Hedgehog signaling pathway<br>(P00025) | 6,10% |
| biological adhesion<br>(GO:0022610) | 3,20% | Axon guidance mediated by netrin<br>(P00009) | 4,10% |
| localization<br>(GO:0051179) | 1,80% | CCKR signaling map (P06959) | 4,10% |

**Table 3.**
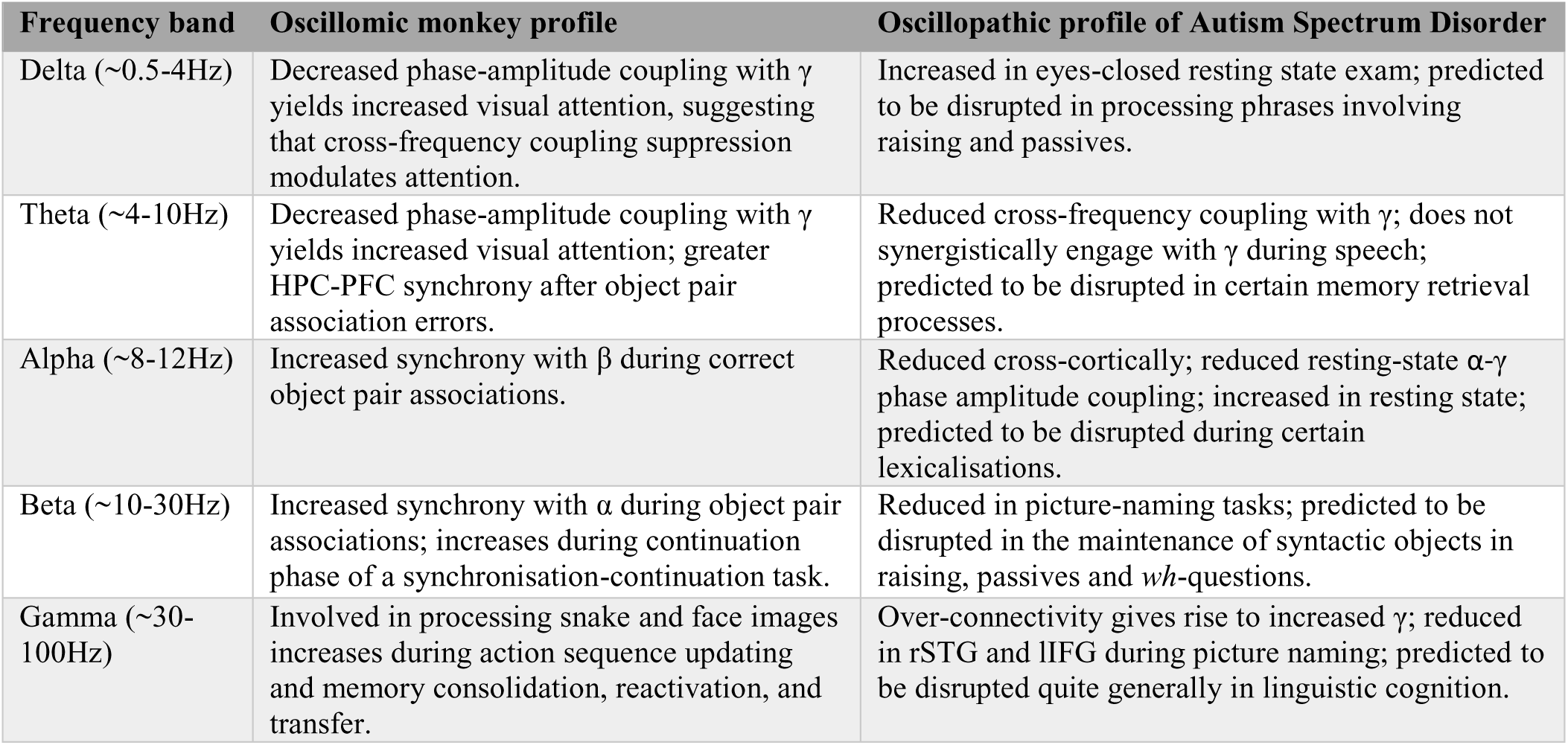
Summary of the patterns of rhythmicity observed in wild primates and the observed oscillomic differences in ASD compared to TD subjects.

### Candidate genes: A functional characterization

Some of Wilkins et al’s (2014) original candidates for the domestication syndrome are candidates for ASD. *KIT* mutations have been found in patients featuring ASD symptoms (Kilsby et al. 2013). KIT is a tyrosine kinase receptor (Kasamatsu et al. 2008), which acts as a key developmental regulator in the NC-derived processes of hematopoiesis, melanogenesis, and gametogenesis (Rothschild et al. 2003). In rats mutations of *Kit* impair hippocampal synaptic potentiation and spatial learning and memory (Katafuchi et al. 2000). Likewise, whole-genome sequencing analyses have identified deleterious variants of *CHD7* in ASD probands (Jiang et al. 2013). *CHD7* is known to be the main candidate for CHARGE syndrome (Vissers et al. 2004, Lalani et al. 2006), mentioned above. Interestingly, CHARGE syndrome also involves microcephaly, face asymmetry, cleft lip/palate, along with variable degrees of intellectual disability (Pisano et al. 2014, Hale et al. 2016 for review). Changes in the expression pattern of *CDH7* can also result in behavioral anomalies resembling the autistic phenotype. Accordingly, *in utero* exposure to heavy metals in mice increases autism-like behavioral phenotypes in adult animals through inducing the hypomethylation of *Chd7* (Hill et al. 2015). *FOXD3*, encoding a transcription factor, is downregulated by DISC1 (Drerup et al. 2009), a robust candidate for schizophrenia that has been also associated to ASD (Williams et al. 2009, Zheng et al. 2011, Kanduri et al. 2016). *FOXD3* maps within one of the present-day human-specific differentially-methylated genomic regions (DMRs) (Gokhman et al. 2014). Interestingly, loss of *Disc1* results in abnormal NCC migration and differentiation (Drerup et al. 2009). Also, DISC1 downregulates *SOX10*, another NC gene, involved in the maintenance of precursor NCC pools, in the timing of NCC migration onset, and in the induction of their differentiation; it is also implicated in oligodendrocyte differentiation (Hattori et al. 2014). In turn, SOX10 interacts with PAX3, another core candidate proposed by Wilkins et al. (2014), and with POU3F2 (Smit et al. 2000). Sequence and copy number variations affecting *POU3F2* have been found in subjects with ASD, and in individuals with different developmental and language delays (Huang et al. 2005, Lin et al. 2011). POU3F2 is a known interactor of *FOXP2*, the renowned “language gene” (Maricic et al. 2013). AMHs bear a derived allele of the binding site which is less efficient in activating transcription than the Neanderthal/Denisovan counterpart (Maricic et al. 2013). Likewise, *POU3F2* has been associated with human accelerated conserved noncoding sequences (haCNSs) (Miller et al. 2014). Also, it interacts with PQBP1, which has been linked to intellectual disability (Wang et al. 2013) and developmental delay and microcephaly (Li et al. 2013). Also *SOX9*, considered a master regulator of craniofacial development and related to several congenital skeletal malformations (Mansour et al. 2002, Gordon et al. 2009, Lee and Saint-Jeannet 2011), is found among the candidates for ASD.

Accordingly, gene and miRNA expression profiling using cell-line derived total RNA has revealed *SOX9* as one of the genes dysregulated in ASD (Ghahramani et al. 2011). As discussed in detail by Benítez-Burraco et al. (submitted) *SOX9* interacts with *BMP2, BMP7, DLX2*_4_and*HES1*. All of them are core components of the network believed important for globularization and language-readiness (reviewed in Boeckx and Benítez-Burraco 2014a). In addition, all of them are involved in NCC development and migration, and in the patterning of NC-derived tissues (Mallo 2001, Gajavelli et al. 2004, Correia et al. 2007, Glejzer et al. 2011, Ishii et al. 2012). *BMP2, BMP7*, and *DLX2* act upstream *SOX9* (Sperber et al. 2008, Li et al. 2013). In turn, *SOX9* mediates the retinoic acid-induced expression of *HES1*, known also to be involved in language function, craniofacial development, and neuron growth and interconnection (reviewed in Boeckx and Benítez-Burraco 2014b). Importantly, retinoic acid also regulates the expression of other genes that are relevant for language, like *FOXP2* (Devanna et al. 2014), or for globularization, like *ASCL1* (see Benítez-Burraco and Boeckx 2014b for details). Retinoic acid has proven to be important for brain plasticity (Luo et al. 2009), and memory and learning processes (Etchamendy et al. 2003, Jiang et al. 2012). Recent whole-exome sequencing analyses have linked retinoic acid regulation pathways to ASD (Moreno-Ramos et al. 2015). In neuronal cells reduced levels of RORA downregulate multiple transcriptional targets that are significantly enriched in biological functions negatively impacted in ASD and which include known ASD-associated genes, like *A2BP1, CYP19A1, ITPR1, NLGN1*, and *NTRK2* (Sarachana and Hu 2013a). *RORA* itself is downregulated in postmortem prefrontal cortex and cerebellum of subjects with ASD (Nguyen et al. 2010). *RORA* is differentially regulated in them by masculine and feminine hormones: Whereas it is under negative feedback regulation by androgens, it is under positive regulation by estrogens (Sarachana et al. 2011, Sarachana and Hu 2013b). In certain regions of the brain this sexually dimorphic expression is also found in several of RORA’s targets and this correlation is much higher in the cortex of males (Hu et al. 2015). Perhaps not surprisingly, synthetic RORa/y agonist improve autistic symptoms in animal models of the disease, particularly, repetitive behavior (Wang et al. 2016).

We wish to highlight two other genes thought to be involved in the changes that brought about modern language that are also candidates for ASD and interact with core candidates for the domestication syndrome as posited by Wilkins et al. The first one is *DLX5*, involved in crucial aspects of NC development (McLarren et al. 2003, Ruest et al. 2003), but also of skull and brain development (Kraus and Lufkin 2006, Wang et al. 2010). Accordingly, it plays a role in thalamic development (Jones and Rubinstein 2004) and contributes to regulate the migration and differentiation of precursors of GABA-expressing neurons in the forebrain (Cobos et al. 2006). *DLX5* is a candidate for ASD (Nakashima et al., 2010), due to an ultraconserved cis-regulatory element (Poitras et al. 2010), which is bound by GTF2I, encoded by one of the genes commonly deleted in Williams-Beuren syndrome (OMIM#194050) and a candidate for ASD too (Malenfant et al. 2012). Additionally, *DLX5* is regulated by MECP2 (Miyano et al. 2008), encoded by the main candidate for Rett syndrome (OMIM#312750), a condition entailing problems for motor coordination, autistic behaviour, and language regression (Uchino et al. 2001, Veenstra-VanderWeele and Cook 2004). Interestingly, Dlx5/6(+/-) mice exhibit abnormal pattern of γ rhythms resulting from alterations in GABAergic interneurons, particularly in fast-spiking interneurons (Cho et al. 2015). In addition, DLX5 interacts with key candidates for language evolution, in particular, with *RUNX2* and *FOXP2* (see Boeckx and Benítez-Burraco 2014a for details). The second one is *NCAM1*, which is also a target of both RUNX2 (Kuhlwilm et al. 2013) and FOXP2 (Konopka et al. 2009). In mice mutations in the gene affect working/episodic-like memory (Bisaz et al. 2013), whereas overexpression of the Ncam1 extracellular proteolytic cleavage fragment impacts on GABAergic innervation, affecting long‐ and short-term potentiation in the prefrontal cortex (Brennaman et al. 2011). *NCAM1* encodes a cell adhesion protein involved in axonal and dendritic growth and synaptic plasticity (Rønn et al. 2000, Hansen et al. 2008). It interacts with VCAM1which is involved in cell adhesion and the control of neurogenesis (Kokovay et al. 2012), and which bears a fixed (D414G) change in AMHs compared to Neanderthals/Denisovans (Pääbo 2014, Table S1). *VCAM1* is upregulated by CLOCK, which plays a key role in the modulation of circadian rhythm (Gao et al. 2014). Together with other circadian-relevant genes *CLOCK* seems to be involved in the psychopathology of ASD cases entailing sleep disturbances (Yang et al. 2016). The circadian modulation of synaptic function has been hypothesised to contribute decisively to ASD (Bourgeron 2007). In turn, *CLOCK* interacts with *RUNX2* and with several other candidates for language-readiness, like *DUSP1*, involved in vocal learning (Doi et al. 2007), and *USF1*, which regulates synaptic plasticity, neuronal survival and differentiation (Tabuchi et al. 2002; Steiger et al. 2004). USF1 binds the promoter of *FMR1* (Kumari and Usdin, 2001), a strong candidate for Fragile-X syndrome (OMIM#300624), which presents with language problems and ASD features (Kaufmann et al. 2004, Smith et al. 2012). The regulatory region of *USF1* shows many fixed or high frequency changes compared to Denisovans (Meyer et al., 2012).

As shown in Table 1, several of the genes important for globularization and language-readiness are involved in NC development and function and some of them are also candidates for ASD. Accordingly, we expect them to contribute to the abnormal domesticated features observed in patients with ASD, and also to their distinctive language profile. Among them we wish mention *CTNNB1, DLX1, DLX6, PAX6*, and *ROBO2*. CTNNB1 is a component of the Wnt/β-catenin signalling pathway, known to be impaired in ASD (Cao et al. 2012, Zhang et al. 2012, Martin et al. 2013). CTNNB1 controls aspects of NC development, from NC induction, lineage decisions, to differentiation (Hari et al. 2012). As noted in Boeckx and Benítez-Burraco (2014b), CTNNB1 is expected to interact with many of the genes highlighted as important for the evolution of language-readiness, specifically with RUNX2 and SLIT2/ROBO1 signals. Regarding DLX1, it is a robust NC marker (Ishii et al. 2012), involved in patterning and morphogenetic processes in NC-derived tissues (Mallo 2001). It also regulates the development of the skull and the brain (Andrews et al. 2003, Jones and Rubenstein 2004). In mice *Dlx1* downregulation results in reduced glutamatergic input to the hippocampus (Jones et al. 2011), as well as in changes in interneuron subtypes and migration patterns in the cortex (Ghanem et al. 2008). *DLX1* is found to be downregulated in ASD (Voienagu et al. 2011, McKinsey et al. 2013). ROBO2 is one of the DLX1 interactors. Slit/Robo signalling regulates early NCC migration (Jia et al. 2005) *ROBO2* is also involved in thalamocortical axons (TCA) development, known to be important for the modulation of cognitive functions (López-Bendito et al. 2007, Marcos-Mondéjar et al. 2012). *ROBO2* is a candidate for ASD (Suda et al. 2011), but also for different types of language disorders, like dyslexia (Fisher et al. 2002) and speech-sound disorder and reading (Stein et al. 2004). It has been related as well to expressive vocabulary growth in the normal population (St Pourcain et al. 2014). Finally, *PAX6* controls the migration of NCCs from the anterior midbrain (Matsuo et al. 1993). *PAX6* is involved as well in the development of the brain (Valverde et al. 2000, Tyas et al. 2003, Caballero et al. 2014). Mutations on *PAX6* have been reported in some forms of ASD (Maekawa et al. 2009), although they also impact in working memory (Bamiou et al. 2007). Alterations of *PAX6* expression in the brain of people with ASD may account for the observed imbalance in excitatory/inhibitory neuronal activity (Kim et al. 2014). And like many of the genes reviewed above, *PAX6* is functionally related to both *FOXP2* and *RUNX2*, and it also targets *POU3F2* (see Benítez-Burraco and Boeckx 2015 for details).

Most of the NCC-genes mentioned here are known to play a key role in the development and patterning of the craniofacial complex, and to be associated to congenital craniofacial defects (Table 1) (see Twigg and Wilkie 2015 for review). Many of these genes are known candidates for ASD, including *DLX5* and *DLX6* (reviewed above), *FGFR2, MSX1, POLR1A*, and *PTCH1*. Both *DLX5* and *DLX6* are indeed required for NC-derived facial morphogenesis (Gitton et al, 2011) FGFRs are among the main craniosynostosis-associated genes. In particular, gain-of-function mutations in *FGFR2* are typically associated to Apert (OMIM#101200) and Crouzon (OMIM#123500) syndromes, while both *FGFR1* and *FGFR2* are found mutated in Pfeiffer syndrome (OMIM#101600) (Lattanzi et al, 2012). All these syndromic craniosynostosic conditions occasionally present with variable degree of ASD-like mental retardation (Morey-Canellas et al. 2003). *MSX1* encodes a transcriptional repressor involved in craniofacial development and shaping (particularly in odontogenesis) (Alappat et al. 2003, Cohen 2000). It is expressed in the NC (Khadka et al. 2006), where it acts as a master regulator of gene expression (Attanasio et al. 2013). Although it has not been associated to ASD, *MSX1* is a direct downstream target of DLX5 during early inner ear formation (Sajan et al. 2011). The gene is also a critical intrinsic dopamine neuron determinant (Andersson et al. 2006) and is found mutated in some patients with Wolf-Hirschhorn syndrome (OMIM#194190), a clinical condition entailing profound mental retardation and craniofacial dysmorphism (Campbell et al. 1989). *POLR1A*, found mutated in acrofacial dysostosis (Cincinnati type, OMIM#616462) involving microcephaly, plays a role in the regulation of NC-derived skeletal precursor cells (Weaver et al. 2015). In some ASD subjects CNVs result in fusion transcripts involving *POLR1A*, although no fusion transcripts have been detected to date (Holt et al. 2012).

Finally, it is worth mentioning that genes encoding primary cilium signaling molecules, such as *SHH, GLI3* and *PTCH1*, are all primarily involved in congenital malformations affecting the midline craniofacial compartment (Brugmann et al. 2010, Rice et al. 2010). Specifically, *PTCH1* is required in the NC-dependent orofacial development and gives rise to orofacial clefting, when mutated (Metzis et al. 2013). Heterozygous mutations of either *SHH* or *PTCH1* are typically found in holoprosencephaly (OMIM#610828, and #236100), a genetically heterogeneous, highly prevalent congenital forebrain anomaly in humans, associated with mental retardation and craniofacial malformations (Mercier et al. 2011, Ming et al. 2002). In addition, a 22-bp deletion in this gene has been found in a girl with ASD and Gorlin syndrome, a complex condition involving macrocrania and hypertelorism (Delbroek et al. 2011).

### Candidate genes: Expression profiles in the ASD brain

If our hypothesis is on the right track, we expect that the genes we highlight here are dysregulated in the brain of people with ASD, particularly in areas important for language processing. Accordingly, we surveyed the Gene Expression Omnibus (GEO) repository (https://www.ncbi.nlm.nih.gov/gds) searching for their expression profiles in the cerebellum and the temporal cortex (but also in the frontal and occipital cortices) in patients with ASD. This should help identify new candidates for ASD in the context of domestication and language-readiness (Table 1). Overall, we could find significant expression values for some of our candidates and learnt that they are up-or downregulated in the brain of autists (Figure 3).

**Figure 3.**
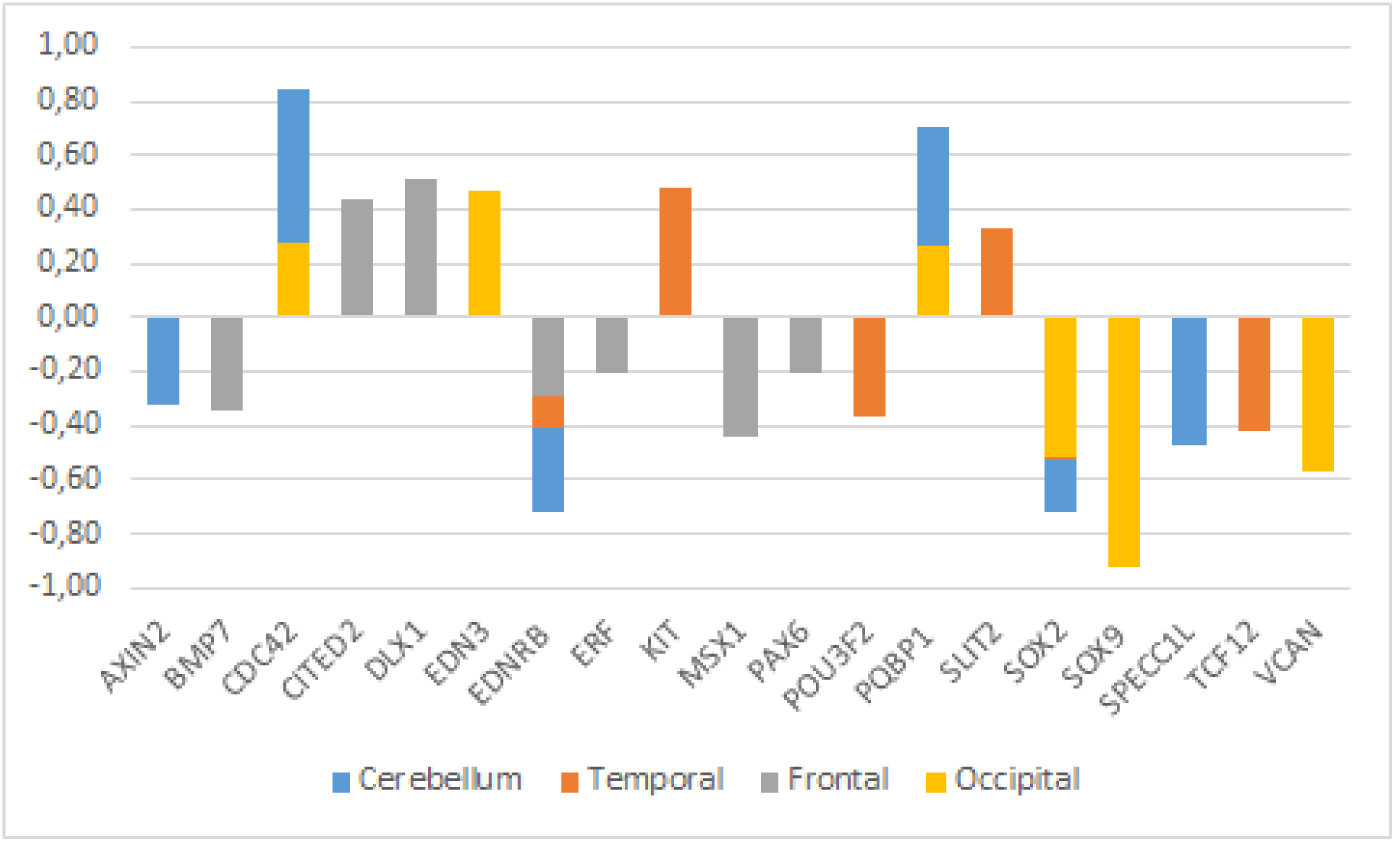
Expression profiles of candidate genes in the ASD brain. Data were gathered from microarray expression datasets available on the GEO datasets: GSE28521 (Voineagu et al. 2011) for the temporal and frontal cortices and GSE38322 (Ginsberg et al. 2012) for the cerebellum and the occipital cortex. Data are shown as log transformation of fold changes (logFC) between patients and corresponding controls. Only genes showing statistically significant (p<0.05) differential expression were considered. Additional details may be found in Supplemental information file.

Among the genes that are significantly downregulated in the cerebellum we found *AXIN2, EDNRB, SOX2, SPECC1L, TCF12*, and *VCAN*, whereas *CDC42*, and *PQBP1* are found upregulated in this region (Figure 3). Although none of them has been associated to ASD, they stroke us as promising candidates for the atypical presentation of the domestication syndrome in ASD. *AXIN2* is expressed in the cranial NC and is needed for NC-derived frontal bone osteogenesis (Yu et al. 2005, Li et al. 2015a). This gene is also expressed as a specific marker for suture stem cells (Maruyama et al. 2016), but also acts as a negative regulator of canonical Wnt pathway, contributing to the stability of CTNNB1 (Li et al. 2015a). Speech alterations are also observed in people with *AXIN2* mutations causing non-syndromic oligodontia (Liu et al. 2015). *EDNRB* encodes a receptor for endothelins, known to be potent vasoactive peptides. Mutations in this gene are associated to increased susceptibility to Hirschsprung disease (OMIM#600155), a neurocristopathy characterized by congenital absence of intrinsic ganglion cells in the enteric nervous plexa (Amiel et al, 2008). Waardenburg syndrome (OMIM#277580), a genetically heterogeneous condition which may involve developmental delay subsidiary to sensorineural hearing loss, has also been associated with mutations in *EDNRB* (Read and Newton 1997). *SOX2*, one of core candidates for domestication (Wilkins et al, 2014), encodes an interactors of the GLI factors as part of the SHH-GLI signaling pathway involved in NCC fate (Oosterveen et al. 2012, Peterson et al. 2012, Oosterveen et al. 2013), but also in the globularization of the AMH skull/brain (see Boeckx et al. submitted for details). SOX2 interacts as well with the BMP signalling (Li et al. 2015b). Interestingly, SOX2 regulates *PQBP1*, highlighted above as one of POU3F2 interactors. *SPECC1L* is found mutated in Opitz G/BBB syndrome (OMIM# #145410) and in facial clefting (Kruszka et al. 2015). This gene functions in NC development (Wilson et al. 2016) and is specifically involved in facial morphogenesis (Saadi et al. 2011).*TCF12* is highly expressed in embryonic precursors of skull/brain structures, including NC-derived head mesenchyme (Uittenbogaard and Chiaramello 2002). *TCF12* directly interacts with *TWIST1*, mutated in Saethre-Chotzen syndrome (OMIM#601622), which features complex craniosynostosys with variable degrees of intellectual disability, including ASD traits (Maliepaard et al, 2014). Indeed, loss-of-function mutations of *TCF12* have been identified in patients with coronal synostosis, which sometimes involves intellectual disability (Sharma et al. 2013, Di Rocco et al. 2014, Paumard-Hernández et al. 2015, Piard et al. 2015). *VCAN* encodes versican-1, a protein that guides migratory NCCs (Dutt et al. 2006) and which shows a fixed N3042D change in AMHs (Pääbo, 2014; Table S1). Finally, *CDC42* controls NC stem cell proliferation (Fuchs et al. 2009). Inactivation of Cdc42 in NCCs causes craniofacial and cardiovascular morphogenesis defects (Liu et al. 2013). As discussed in detail in Boeckx and Benítez-Burraco (2014b) *CDC42* is an important gene regarding language evolution, because of its functional connections with core candidates for globularization and the externalization of language (including *FOXP2, RUNX2, SLIT2*, and *ROBO1*), with genes related to language disorders (like *ITGB4, ARHGAP32, ANAPC10* and *CDC42EP4*).

With regards to the temporal cortex, we found that *EDNRB, POU3F2, SOX2, VCAN*, and *TCF12* are downregulated in subjects with ASD, whereas *KIT, PQBP1*, and *SLIT2* are upregulated in them (figure 3). As noted above, *POU3F2, SOX2, KIT* are known candidates for ASD. We have already reviewed all these genes. Concerning the frontal cortex we found that *BMP7, EDNRB, PAX6, ERF*, and *MSX1* are significantly downregulated, whereas *CITED2* and *DLX1* are significantly upregulated (Figure 3). As noted in table 1, *DLX1, PAX6*, and*MSX1* have been previously associated to ASD. Most of these genes have been already reviewed. *BMP7* is a NCC gene involved in regulation of osteogenesis (Cheng et al. 2003) and in skull and brain development (Yuge et al. 2011, Segklia et al. 2012). *BMP7* is also closely related to some of the core candidates for globularization and language-readiness, like *BMP2, DLX1, DLX2*, and *RUNX2*. BMP7 is predicted (according to String 10) to interact with SOX2 via NOG, involved in dopamine neuron production and an inhibitor of BMP signalling (Chiba et al. 2008). Developmental delay and learning disabilities are commonly observed in people with mutations in *BMP7* (Wyatt et al. 2010). *ERF* encodes a member of the ETS family of transcription factors, expressed in migratory cells, including NCCs (Paratore et al. 2002). *ERF* haploinsufficiency gives rise to either coronal or multisuture synostosis, midface hypoplasia, often associated with behavioural and learning difficulties (Twigg et al. 2013). Concerning *CITED2*, this is a functional partner of both FOXP2 and RUNX2 (Luo et al. 2005, Vernes et al. 2011, Nelson et al. 2013), two important genes for the emergence of modern language. *CITED2* is also involved in craniofacial development (Bhattacherjee et al. 2009) and in the establishment of left-right axis through interactions with the BMP signaling and Nodal (Preis et al., 2006, Lopes Floro et al. 2011). Interestingly, 99% of AMHs bear a highly disruptive intergenic change near *CITED2* compared to Altai Neanderthals and Denisovans (Prüfer et al. 2014).

Finally, regarding the occipital cortex, we found that *EDN3, CDC42*, and *PQBP1* are significantly upregulated in ASD, whereas *SOX2, SOX9*, and *VCAN* are downregulated. We have already considered all these genes, with the exception of *EDN3*. This gene encodes an endothelium-derived vasoactive peptide which binds the product of *EDNRB*, playing a key role in the development of neural crest-derived cell lineages, such as melanocytes and enteric neurons. Although *EDN3* is a candidate for Waardenburg syndrome and Hirschsprung disease, the gene has been found significantly dysregulated in children with ASD (Glatt et al. 2011).

As shown in Figure 3, genes that exhibit significant changes in their expression levels in the ASD brain are consistently down-or upregulated across all the regions under analysis. Considering their functions and the phenotypes resulting from their mutation, the most promising of these genes are *CDC42* and *PQBP1* (which are upregulated) and *EDNRB* and *SOX2* (which are found downregulated). As noted above, to date none of them have been associated to ASD, but they emerge as reasonably involved in the pathogenesis of this condition.

Noteworthy age-related differences in the intrinsic functional connectivity of the brain are observed in ASD: adult patients show reduced connectivity while children tend to exhibit an increased connectivity (Uddin et al. 2013). Therefore, we have further analysed ASD brain expression data in an age-matched fashion. Due to the available sample characteristics, only expression data obtained from the cerebellum could be analysed (see Supplemental information file for further details). The reduction of the age-related bias in the patients-versus-controls comparison, enabled finding a higher number of statistically significant dysregulated genes in the ASD brain. Accordingly, we found that in the cerebellum of ASD children (below 11 years old), *CDC42, MSX1, MSX2, NODAL, PQBP1*, and *SLIT2* were downregulated, whereas, *AXIN2, CHD7, CITED2, EDNRB, FGF8, NCAM1, PAX7, PTCH1, RET, ROBO1, SOX2, SPECC1L, TCF12, VCAN*, and *ZIC1* were upregulated. In turn, in the cerebellum of adult patients (aged 22-to-60 years) we found that *CDC42, CTNNB1, DLX1, EDNRA, EDNRB, HES1, KIT, MAGOH, MITF, NCAM1, NTN1, POU3F2, PQBP1, PTCH1, RET, ROBO1, SATB2, SOX2, SPECC1L, TCF12, VCAN*, and *ZIC1* are downregulated, whereas only *AXIN2, CTNNB1, DLX1, EDN1*, and *MSX1* are upregulated. Overall, we concluded that nearly one third of the candidates for domestication are dysregulated in the cerebellum of people with ASD, and that more than a half are specifically dysregulated in the cerebellum of either children and/or adults with this condition. Genes that are dysregulated in both children and adults with ASD can be regarded as significant contributors to the atypical presentation of the domestication syndrome in this condition. This list encompasses 14 genes: *AXIN2, CDC42, EDNRB, MSX1, NCAM1, PQBP1, PTCH1, RET, ROBO1, SOX2, SPECC1L, TCF12, VCAN*, and *ZIC1*. Interestingly, they exhibit opposite expression profiles in children and adults with ASD (figure 4). Most of these genes have been already discussed here. In addition,*RET*, encoding a cadherin that plays a crucial role in NC development, is a candidate for Hirschsprung disease (OMIM# 142623; Edery et al. 1994) and is found to be differentially expressed after RUNX2 transfection in neuroblastoma cells (Kuhlwim et al. 2013). *RET* is downstream *ASCL1* (another candidate for Hirschsprung disease) in noradrenergic brain stem neurons important for respiratory rhythm modulation (Dauger et al. 2011). Likewise, *ZIC1* is needed for NC development (along with *PAX3*) and plays a key role in craniofacial development (Milet et al, 2013, Plouhinec et al, 2014). Mutations in *ZIC1* result in severe coronal synostosis associated with learning difficulties (Twigg et al. 2015).

**Figure 4.**
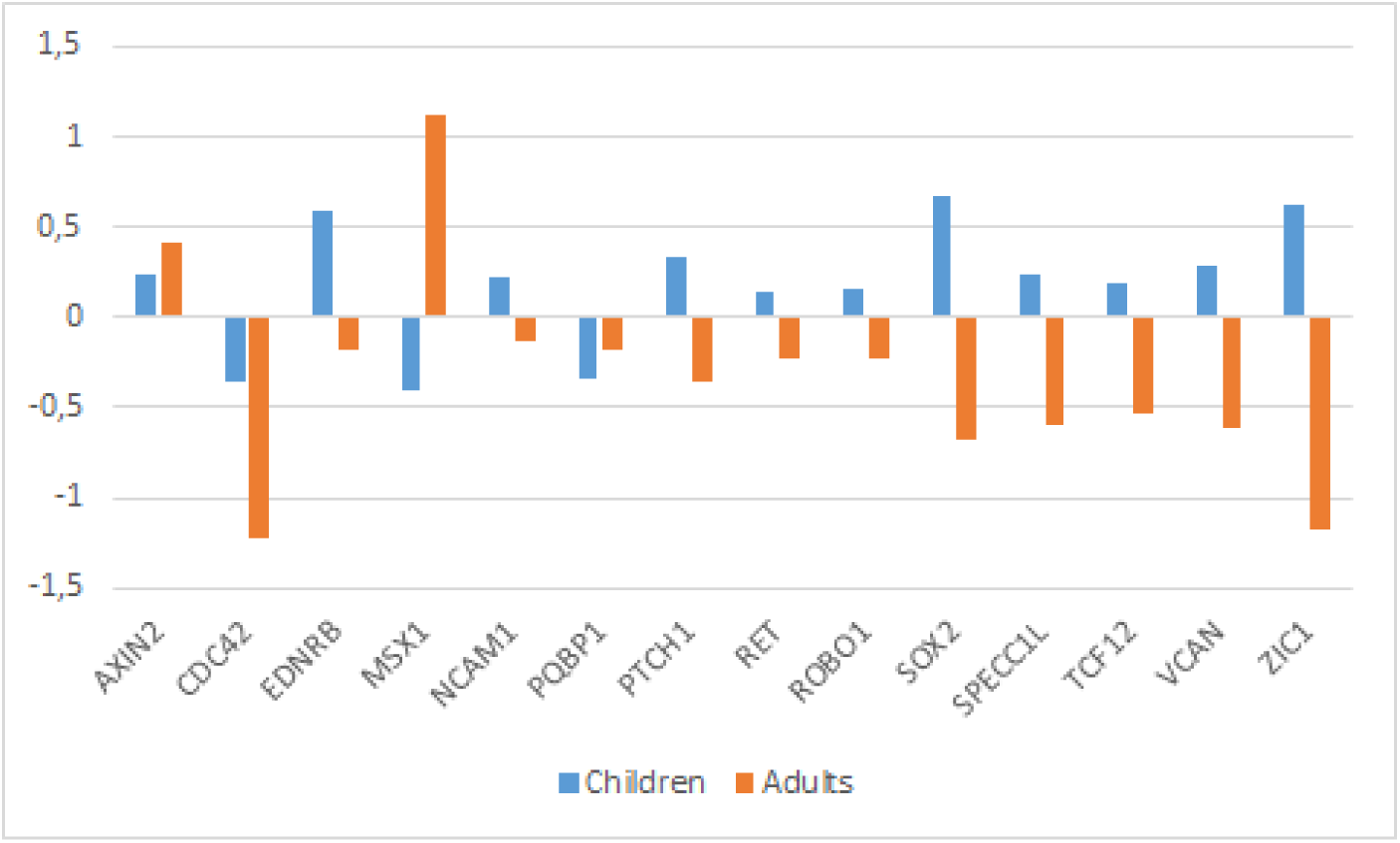
Expression profiles of candidates genes in the cerebellum of children and adults with ASD. Expression data were obtained from the microarray expression dataset GSE38322 (Ginsberg et al. 2012) available on the Gene Expression Omnibus database (GEO datasets, http://www.ncbi.nlm.nih.gov/gds). Data are shown as log transformation of fold changes (logFC) between patients and corresponding controls. Only genes showing statistically significant (p<0.05) differential expression were considered. Additional details may be found in Supplemental information file.

In the last section of the paper we attempt refining our characterization of ASD as an atypically-domesticated phenotype. In doing so, we will focus on brain function, with a special emphasis on brain oscillations. Accordingly, we will compare the oscillopathic profile of people with ASD during language processing with the oscillatory signature of the TD population and that of non-domesticated primates. Additionally, we will examine the expression profile in the primate brain of the candidate genes we highlight here.

## 4. ASD and wild primates: from brain oscillation to gene expression patterns

### ASD and primate oscillomes

As we have discussed in detail in Benítez-Burraco and Murphy (2016) language impairment in ASD (as ASD itself) can be satisfactorily characterised as an oscillopathic condition. With cognitive disorders exhibiting disorder-specific abnormal oscillatory profiles, it is also noteworthy that species-specific oscillatory patterns seemingly emerge as slight variations within the network constellation that constitutes a universal brain syntax (Buzsáki and Watson 2012 Buzsáki et al. 2013).

The differences in brain capacity between domesticated and non-domesticated animals (Wilkins et al. 2014) would be predicted to give rise to a corresponding alteration in oscillatory properties (i.e. features of neural oscillations which form part of an individual’s ‘oscillome’, as it is termed in Murphy and Benítez-Burraco 2016 and Murphy 2016). Although we feel that current knowledge is too scarce to permit any reasonable linking hypotheses between the primate and ASD oscillomes, we would like to briefly sketch out a possible route to increasing our understanding of the neural signature of domestication (and failed domestication itineraries).

Call vocalizations have been found not to be impaired when the homologue of Broca’s region in the monkey brain is lesioned, which suggests that other area (like the limbic system and brainstem) are involved (Sage 2006). However, macaques appear to share similar call comprehension substrates with human language comprehension in the left posterior temporal gyrus (Heffer & Heffner 1986). It would be of interest, for instance, to compare the rhythmic properties of this region of the TD brain with those of the primate brain to see if any particular activity (e.g. coupling and synchronization) is marked in humans. This would also yield insights into how the primate call comprehension system ‘interfaces’ with other cognitive systems (given the appropriate experimental environment), and would also permit the exploration of similar interface properties of human language comprehension, which requires the transfer of information to two interfaces (Figure 5).

**Figure 5.**
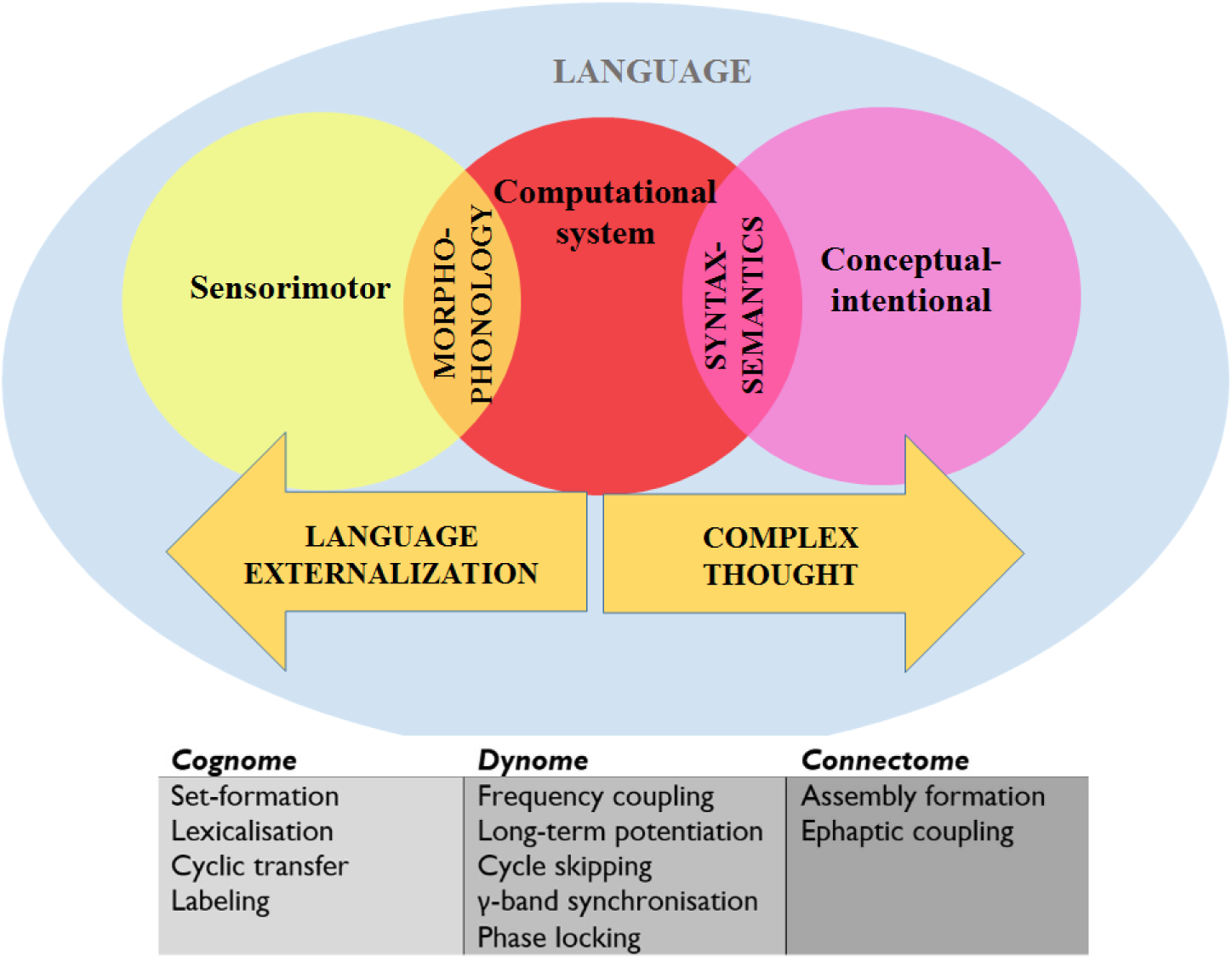
A schematic view of language representing the systems and interfaces of interest and the levels of analysis as discussed in the paper. ‘Cognome’ refers to the operations available to the human nervous system (Poeppel, 2012) and ‘dynome’ refers to brain dynamics (Kopell et al., 2014) (see Murphy and Benítez-Burraco 2016 for details).

Discounting work on evoked potentials (which is itself fairly modest), there are currently only a handful of empirical studies of the monkey oscillome. Brincat and Miller (2015), for instance, discovered functional differences and frequency-specific interactions between the Rhesus hippocampus (HPC) and prefrontal cortex (PFC) during object pair association learning γ synchrony was found to be greater after errors and decreased after learning; correct associations increased β-α synchrony, which was also greater in the HPC-PFC direction. Esghaei et al. (2015) also suggested that the macaque visual cortex employs phase-amplitude coupling to regulate inter-neuronal correlations, and so the potential for generic oscillomic processes to yield insights into cognitive (dys)function seems apparent. During the internally monitored continuation phase of a synchronisation-continuation task, β also appears to increase in Rhesus monkeys (Bartolo & Merchant 2015), suggesting that - as in humans (Murphy 2015a) - β is responsible for maintaining the existing cognitive set in memory. β is also involved in ‘the attentive state and external cues as opposed to detailed muscle activities’ in Japanese monkeys (*Macaca fuscata*) (Watanabe et al. 2015). Finally, pulvinar γ is involved in feedforward processing for snake images, and also in cortico-pulvinar-cortical integration for face images (Le et al. 2016), while Ramirez-Villegas et al.’s (2015) study of the macaque hippocampal CA3-CA1 network pointed to the role of γ in memory reactivation, transfer and consolidation.

Currently, there are no studies comparing the oscillome of domesticated and non-domesticated primates, but our prediction would be that non-domesticated primates display a degree of oscillomic difference with domesticated primates comparable to the difference between TD individuals and people with ASD. Table 2 summarises existing knowledge of the ASD and primate oscillome during a range of cognitive tasks, and it is hard to find any correlations or connections between the two. However, we feel that comparatively exploring these oscillomes will permit a greater understanding of the atavistic neural oscillations of the non-domesticated human and primate brains. Future research should also seek to compare the oscillomes of domesticated and non-domesticated primates in an effort to investigate neural signatures of domestication.

### Candidate genes: Expression profiles in the primate brain

If our hypothesis turns to be on the right track, we further expect that the genes that we have found dysregulated in the brain of people with ASD show similar expression profiles in conditions where normal socialization failed to occur. Because feral children are scarce and not easily available we examined the expression profiles of these genes in wild primates (chimps). In particular, we selected available gene expression profiles obtained from chimp brain areas that are known to be involved in language processing in humans (the cerebellum, the temporal cortex, and the frontal cortex), as we did for people with ASD. We learnt that most of the genes that we had previously found differentially expressed in the ASD brain data exhibit the same expression pattern in the chimp brain, including *EDNRB* (in the cerebellum), *BMP7, DLX1, EDNRB, MSX1*, and *PAX6*(in the frontal cortex), and *VCAN* (in the temporal cortex) (figure 6).

**Figure 6.**
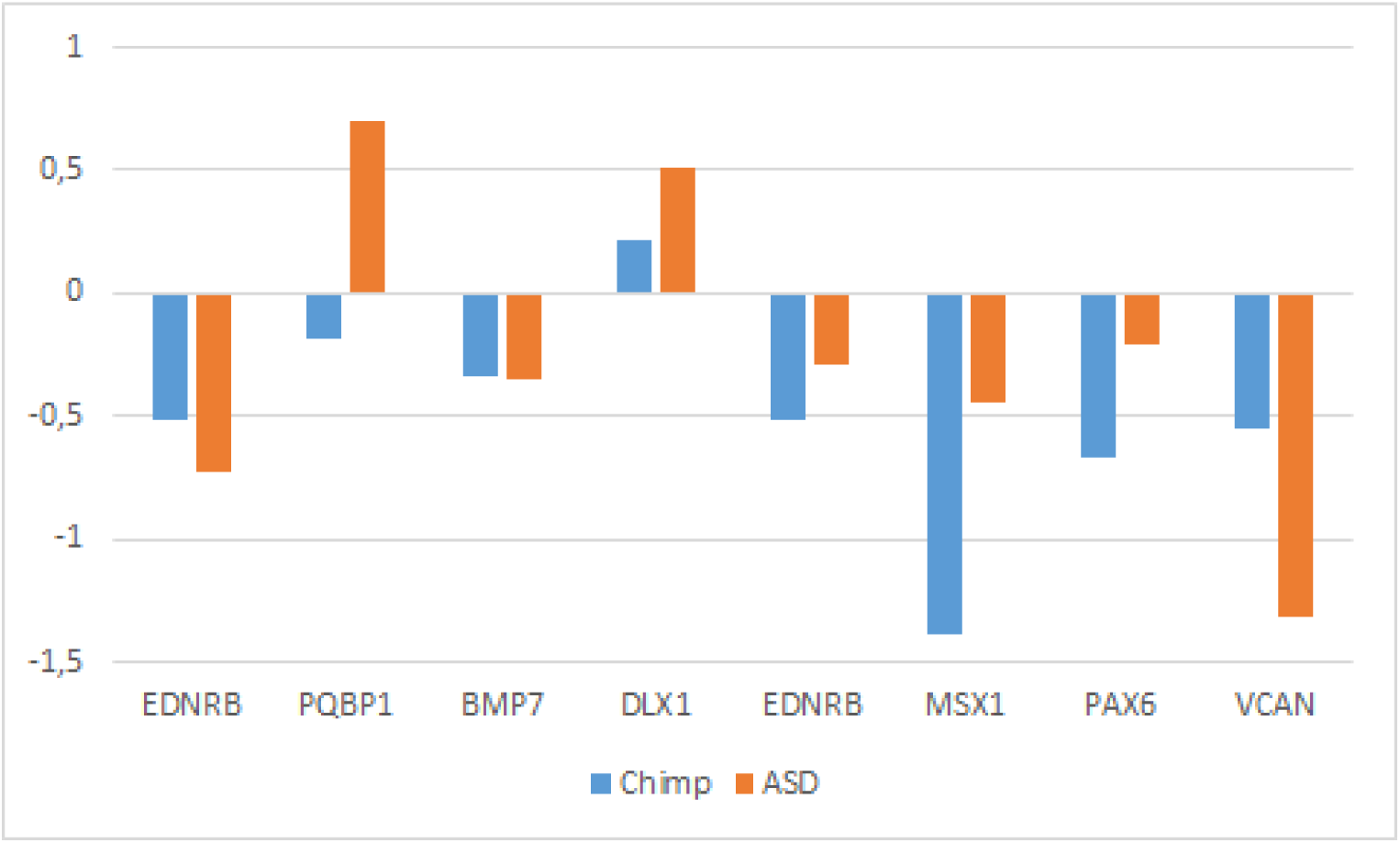
Comparative expression profiles in chimpanzees and subjects with ASD of candidate genes. Data were obtained from microarray expression datasets available on the Gene Expression Omnibus database (GEO datasets, www.ncbi.nlm.nih.gov/gds): GSE28521 (Voineagu et al, 2011) for the temporal and frontal cortices, and GSE38322 (Ginsberg et al. 2012) for the cerebellum of subjects with ASD; and GSE22569 (Somel et al. 2011, Liu et al. 2012) for the cerebellum, GSE18142 (Konopka et al. 2009) for the frontal cortex, and GSE7540 (Cáceres et al. 2003) for the temporal cortex of chimps. Data are shown as log transformation of fold changes (logFC) between patients and corresponding controls. Only genes showing statistically significant (p<0.05) differential expression were considered. Additional details may be found in Supplemental information file. Note that the plot is intended to display the overall trend of gene expression, given that the relative expression values (i.e. logFC) were obtained from comparative analyses performed on different datasets (with inherent different designs, samples, and batches).

## 5. Conclusions

Socialization is a crucial step needed for the achievement of many cognitive abilities that are a signature of the human condition. Language is one of the most prominent of such abilities. Several high prevalent pathological conditions impact on human-specific cognitive abilities, including schizophrenia and ASD. In ASD, social abilities are seriously compromised, but other core cognitive skills, including language, also exhibit differences with the non-affected population. ASD is a multifactorial condition, with wide clinical and genetic heterogeneity. It is still not clear how ASD features emerge from genomic and/or environmental cues during development. In this paper we have focused on language deficits in ASD, although we expect that the lessons we draw here contribute to shedding light on the whole profile of this cognitive condition. In doing so we have adopted and evolutionary perspective, because of the robust link that exists between (abnormal) development and evolution. As we have shown, domesticated traits are absent or are attenuated in people with ASD, and genes that we believe important for the (self)domestication of our species and the evolution of our distinctive cognitive abilities (including language) show abnormal expression patterns in the brains of people with autism. What is more: abnormalities can be traced to the time window when crucial brain rewiring occurs during language acquisition and when changes in the normal configuration of the brain occurs in children with ASD. Additionally, some features of the ASD phenotype can be found (or expected to be found) in wild primates. On the whole, we think that our approach can help illuminate the aetiology of ASD primarily because provides robust links between the genome and the environment, and between development and evolution, in line with the current evo-devo approaches to cognitive diseases (see Benítez-Burraco 2016a for review). In this sense, the putative involvement of the neural crest in the aetiopathogenesis of ASD emerges as a promising avenue for future research. At the same time, we expect that our approach with help illuminate the evolutionary history of our language-readiness: Our results support the view that language evolution benefitted from a favourable social context that may have resulted from our (self)-domestication.

## Acknowledgements

Preparation of this work was supported in part by funds from the Spanish Ministry of Economy and Competitiveness (grant numbers FFI2014-61888-EXP and FFI-2013-43823-P to ABB), in part by "Linea D1-2015" intramural funds from Università Cattolica S. Cuore (to WL), and in part by an Economic and Social Research Council scholarship (1474910) (to EM).

